# Conservation of specificity in two low-specificity protein

**DOI:** 10.1101/207324

**Authors:** Lucas C. Wheeler, Jeremy A. Anderson, Anneliese J. Morrison, Caitlyn E. Wong, Michael J. Harms

**Affiliations:** Department of Chemistry and Biochemistry, University of Oregon, Eugene, OR, USA.; Institute of Molecular Biology, University of Oregon, Eugene, OR, USA

## Abstract

S100 proteins bind linear peptide regions of target proteins and modulate their activity. The peptide binding interface, however, has remarkably low specificity and can interact with many target peptides. It is not clear if the interface discriminates targets in a biological context, or whether biological specificity is achieved exclusively through external factors such as subcellular localization. To discriminate these possibilities, we used an evolutionary biochemical approach to trace the evolution of paralogs S100A5 and S100A6. We first used isothermal titration calorimetry to study the binding of a collection of peptides with diverse sequence, hydrophobicity, and charge to human S100A5 and S100A6. These proteins bound distinct, but overlapping, sets of peptide targets. We then studied the peptide binding properties of S100A5 and S100A6 orthologs sampled from across five representative amniote species. We found that the pattern of binding specificity was conserved along all lineages, for the last 320 million years, despite the low specificity of each protein. We next used Ancestral Sequence Reconstruction to determine the binding specificity of the last common ancestor of the paralogs. We found the ancestor bound the whole set of peptides bound by modern S100A5 and S100A6 proteins, suggesting that paralog specificity evolved by subfunctionalization. To rule out the possibility that specificity is conserved because it is difficult to modify, we identified a single historical mutation that, when reverted in human S100A5, gave it the ability to bind an S100A6-specific peptide. These results indicate that there are strong evolutionary constraints on peptide binding specificity, and that, despite being able to bind a large number of targets, the specificity of S100 peptide interfaces is indeed important for the biology of these proteins.

## Introduction

Many proteins have low specificity interfaces that can interact with a wide variety of targets (1–11). Such interfaces are difficult to dissect. Crucially, it is not obvious that their specificity is biologically meaningful: maybe such proteins are essentially indiscriminate, and biological specificity is encoded by external factors such as subcellular localization or expression pattern (3, 12, 13).

An evolutionary perspective allows us to probe whether specificity is, indeed, an important aspect of these interfaces (14). If there are functional and evolutionary constraints on binding partners, we would expect conservation of binding specificity similar to that observed for high-specificity protein families (15, 16). In contrast, if specificity is unimportant, we would expect it to fluctuate randomly over evolutionary time. Further, previous work on the evolution of specificity has revealed common patterns for the evolution of specificity (17–19), including partitioning of ancestral binding partners among descendant lineages (20–23) and transitions through more promiscuous intermediates (10, 24, 25). If low-specificity proteins exhibit similar patterns, it is strong evidence that the low specificity interface has conserved binding properties, and that the interface makes a meaningful contribution to biological specificity.

S100 proteins are an important group of low-specificity proteins (26, 27). Members of the family act as metal sensors (28), pro-inflammatory signals (29–32), and antimicrobial peptides (33). Most S100s bind to linear peptide regions of target proteins via a short hydrophobic interface exposed on *Ca*^2+^-binding (Fig 1A). S100s recognize extremely diverse protein targets (27, 34, 35). No simple sequence motif for discriminating binders from non-binders has yet been defined. The breadth of targets is much more extreme than other low-specificity proteins such as kinases and some hub proteins, which recognize well-defined, but degenerate, sequence motifs (1,3, 6, 10, 11).

**Fig 1.**
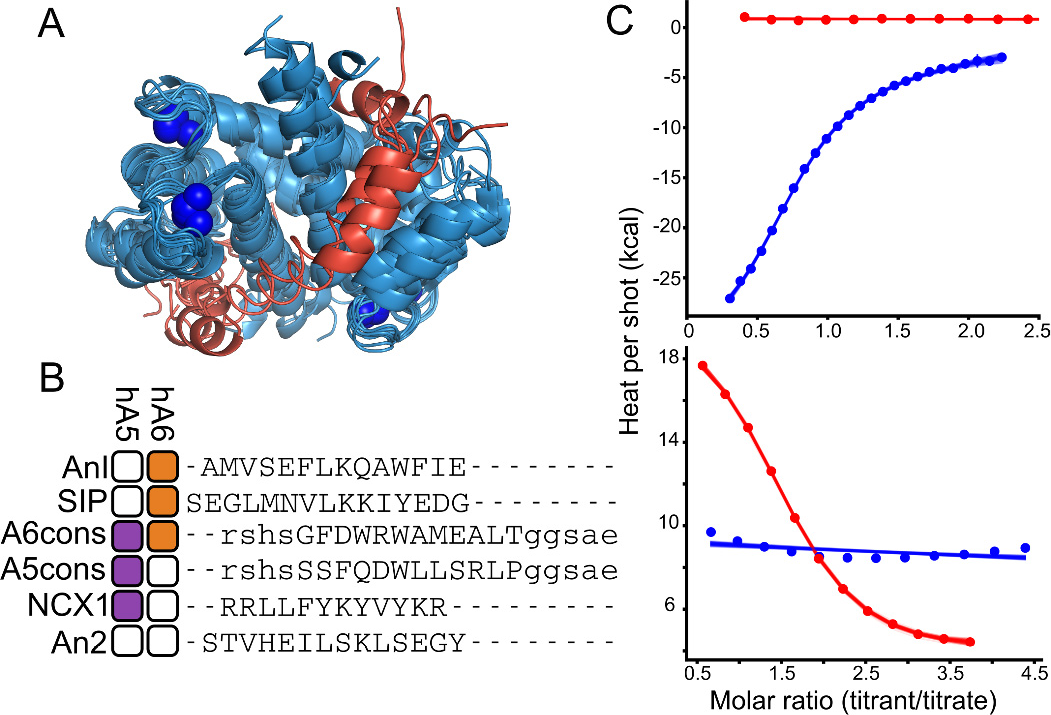
Human S100A5 and S100A6 exhibit peptide binding specificity. A) Published structures of S100 family members bound to both *Ca*^2+^ and peptide targets at the canonical hydrophobic interface (PDB: 3IQQ, 1QLS, 3RM1, 2KRF, 4ETO, 2KBM, 1MWN, 3ZWH). Structures are aligned to the *Ca*^2+^-bound structure of human S100A5 (2KAY). Peptides are shown in red. Blue spheres are *Ca*^2+^ ions. B)Binding specificity of hA5 and hA6. Boxes indicate whether the peptide binds to hA5 (purple) and/or hA6 (orange). If peptide does not bind by ITC (*K*_*D*_ ⪆ 100 *μM*), the box is white. Peptide names are indicated on the left. Peptide sequences, aligned using MUSCLE (75), are shown on the right. Solubilizing flanks, which contribute minimally to binding (Table S1), are shown in lowercase letters. Annexin 1 (An1) and Annexin 2 (An2) binding measurements are from a published study (35). C) ITC heats for the titration of NCX1 (blue) and SIP (red) peptides onto hA5 (top) and hA6 (bottom). Points are integrated heats extracted from each shot. Lines are 100 different fit solutions drawn from the fit posterior probability distributions. For the hA5/NCX1 and hA6/SIP curves, we used a single-site binding model. For hA5/SIPand hA6/NCX1, we used a blank dilution model these fits are in Table S2-S5.

We set out to determine whether there was conserved specificity for two S100 paralogs, S100A5 and S100A6. These proteins arose by gene duplication in the amniote ancestor ≈ 320 million years ago (36, 37). S100A6 regulates the cell cycle and cellular motility in response to stress (38). It binds to many targets including p53 (39, 40), RAGE (31), Annexin A1 (35), and Siah-interacting protein (41). A crystal structure of human S100A6 bound to a fragment of Siah-interacting protein revealed that peptides bind via the canonical hydrophobic interface shared by most S100 proteins (41). The biology of S100A5 is less well understood. It binds both RAGE (31, 32) and a fragment of the protein NCX1 (42) at the canonical binding site. It is highly expressed in mammalian olfactory tissues (43–45), but its specific targets and their biological roles are not well understood.

Using a combination of *in vitro* biochemistry and molecular phylogenetics, we addressed three key questions regarding the evolution of specificity in S100A5 and S100A6. First: do the two human proteins exhibit specificity relative to one another? Second: is the set of binding partners recognized by each protein fixed over time, or does the set of partners fluctuate? And, third: do we see similar patterns of specificity change after gene duplication for these low-specificity proteins compared to high-specificity proteins? Unsurprisingly, we find that S100A5 and S100A6 both bind to a wide variety of diverse peptides. Surprisingly, we find that the set of partners, despite being diverse, has been conserved over hundreds of millions of years. Further, we observe a pattern of subfunctionalization for these low-specificity proteins that is identical to that observed in high-specificity proteins. This suggests that these low-specificity interfaces are indeed constrained to maintain a specific—if large—set of binding targets.

## Results

### Human S100A5 and S100A6 interact with diverse peptides at the same binding site

We first systematically compared the binding specificity of human S100A5 (hA5) relative to human S100A6 (hA6) for a collection of six peptides (Fig 1B). Peptide targets have been reported for both hA5 and hA6 (31, 32, 35, 39–42), but only two targets have been directly compared between paralogs. Using Isothermal Titration Calorimetry (ITC), Streicher and colleagues found that a peptide fragment of An-nexin 1 bound to hA6 but not hA5, and a peptide fragment of Annexin 2 bound to neither (35) (Fig 1B). To better quantify the relative specificity of these proteins, we used ITC to measure the binding of two additional peptides to recombinant hA5 and hA6. The first was a peptide from Siah-interacting protein (SIP) previously reported to bind to hA6 (41). We found that this peptide bound to hA6 with a *K*_*D*_ of 20 *μM*, but did not bind hA5 (Fig 1B, C). The second was a 12 amino acid fragment of the protein NCX1 that was reported to bind to hA5 (42). We found that this peptide bound with to hA5 with a *K*_*D*_ of 20 *μM*, but did not bind hA6 (Fig 1B, C).

To further characterize the specificity of the interface, we used phage display to identify two additional peptides that bound to each protein. We panned a commercial library of random 12-mer peptides fused to M13 phage with either hA5 or hA6. Phage enrichment was strictly dependent on *Ca*^2+^ (Fig S1). Three sequential rounds of binding and amplification with either hA5 or hA6 led to enrichment of the “A5cons” and “A6cons” peptides (Fig 2B, Fig S1). We then used ITC to measure binding of these peptides to hA5 and hA6. To ensure solubility, we added polar N- and C- terminal flanks before characterizing binding. A5cons bound to both hA5 and hA6 (Fig 1C). In contrast, A6cons, bound hA6 but not hA5 (Fig 1C). To verify that binding was driven by the central region, we re-measured binding in the presence and absence of different versions of the flanks (Table S1).

**Fig 2.**
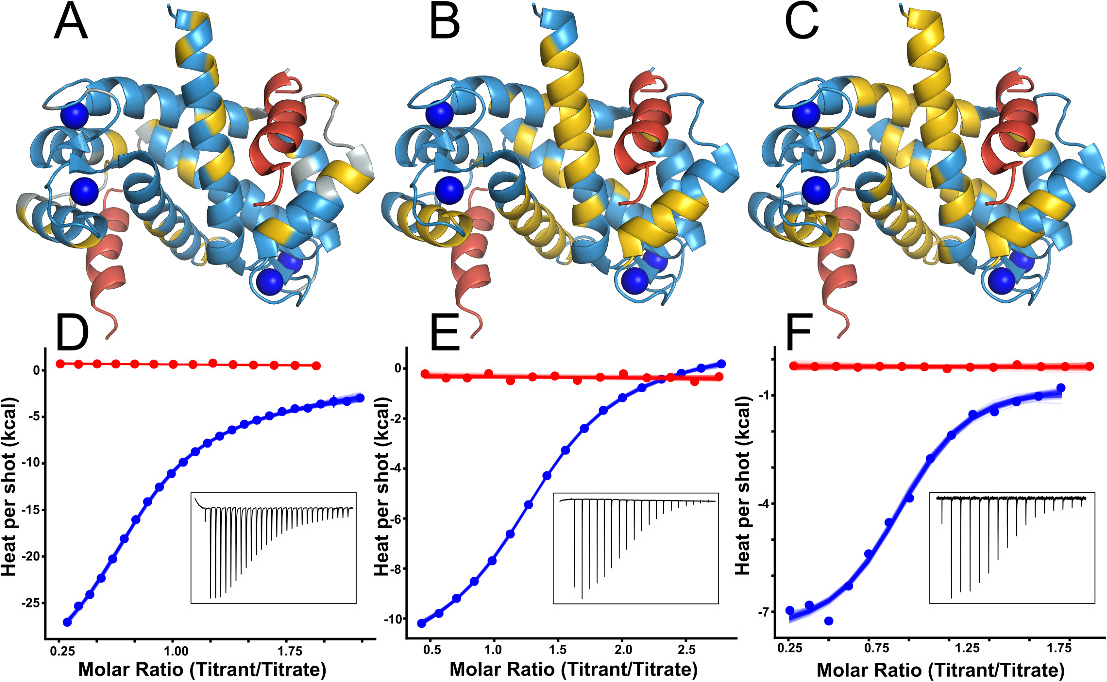
Diverse peptides bind at the human S100A5 peptide interface. Structures show NMR data mapped onto the structure of *Ca*^2+^-bound hA5 (2KAY (46)). To indicate the expected peptide binding location, we aligned a structure of hA6 in complex with the SIP peptide (2JTT (41)) to the hA5 structure, and then displayed the SIP peptide in red. Panels A-C show binding for NCX1, A5cons, and A6cons respectively. In panel A, yellow residues are those noted as responsive to NCX1 binding in (42). In panels B and C, yellow residues are those whose ^1^*H* –^15^ *N* TROSY-HSQC peaks could not be identified in the peptide-bound spectrum because the peaks either shifted or broadened. Panels D-E show ITC data for binding of the peptides above in the presence of 2 mM *Ca*^2+^ (blue) or 2 mM EDTA (red). Points are integrated heats extracted from each shot. Lines are 100 different fit solutions drawn from the fit posterior probability distributions. For the *Ca*^2+^ curves, we used a single-site binding model. For the EDTA curves, we used a blank dilution model. Insets show raw ITC power traces for the *Ca*^2+^ binding curves. Thermodynamic parameters for these fits are in Table S2-S5.

The peptides that bind to hA5 and hA6 are diverse in sequence, hydrophobicity, and charge (Fig 1B). One explanation for this diversity could be that the peptides bind at different interfaces on the protein. To test for this possibility, we used NMR to identify residues whose chemical environment changed on binding of peptide. We first verified the published assignments for hA5 using a 3D NOESY-TROSY experiment (46). We then collected ^1^*H* –^15^ *N* TROSY-HSQC NMR spectra of *Ca*^2+^-bound protein in the presence of either the A5cons or A6cons peptide. By comparing the bound and unbound spectra, we could identify peaks whose location shifted dramatically or that broadened due to exchange. In addition to our own work, we also included previously reported experiments probing the hA5/NCX1 peptide interaction in the analysis (42). For all three peptides, we observed a consistent pattern of perturbations in helices 3 and 4 and, to a lesser extent, helix 1 upon peptide binding (Fig 2A-C). These results suggest that all three peptides bind at the canonical interface. In addition to this spectroscopic evidence, binding of all of these peptides was strictly dependent on the presence of *Ca*^2+^ (Fig 2D-F)—consistent with binding at the interface exposed on *Ca*^2+^ binding (46).

### The S100A5 and S100A6 clades exhibit conserved binding specificity

Although hA5 and hA6 exhibit distinct specificity relative to one another (Fig 1B). This could either result from functional constraints or, alternatively, simply be chance. These possibilities can be distinguished with an evolutionary perspective. If specificity at the interface is functionally important, we would expect conserved specificity between paralogs; if it is unimportant, we would expect it to fluctuate over evolutionary time. We therefore set out to study the evolution of the differences in peptide binding between the human proteins.

We first constructed a maximum-likelihood phylogeny of the clade containing S100A2, S100A3, S100A4, S100A5, and S100A6 (Fig 3A). We built the tree using the EX/EHO+Г_8_ evolutionary model (47), which uses different evolutionary models for sites in different structural classes. As expected from previous phylogenetic and syntenic analyses (37, 48), S100A5 and S100A6 were paralogs that arose by gene duplication in the amniote ancestor, with S100A2, S100A3, and S100A4 forming a closely-related out group (Fig 3A). To set our expectation for conservation of specificity, we then calculated the conservation of residues at the binding site across S100A5 and S100A6 homologs. Fig 3B and C show the relative conservation of residues on hA5 (Fig 3B) and hA6 (Fig 3C). Taken as a whole, the peptide binding region does not exhibit higher conservation than other regions in the protein. We therefore predicted substantial variability in the peptide binding specificity across S100A5 and S100A6 orthologs.

**Fig 3.**
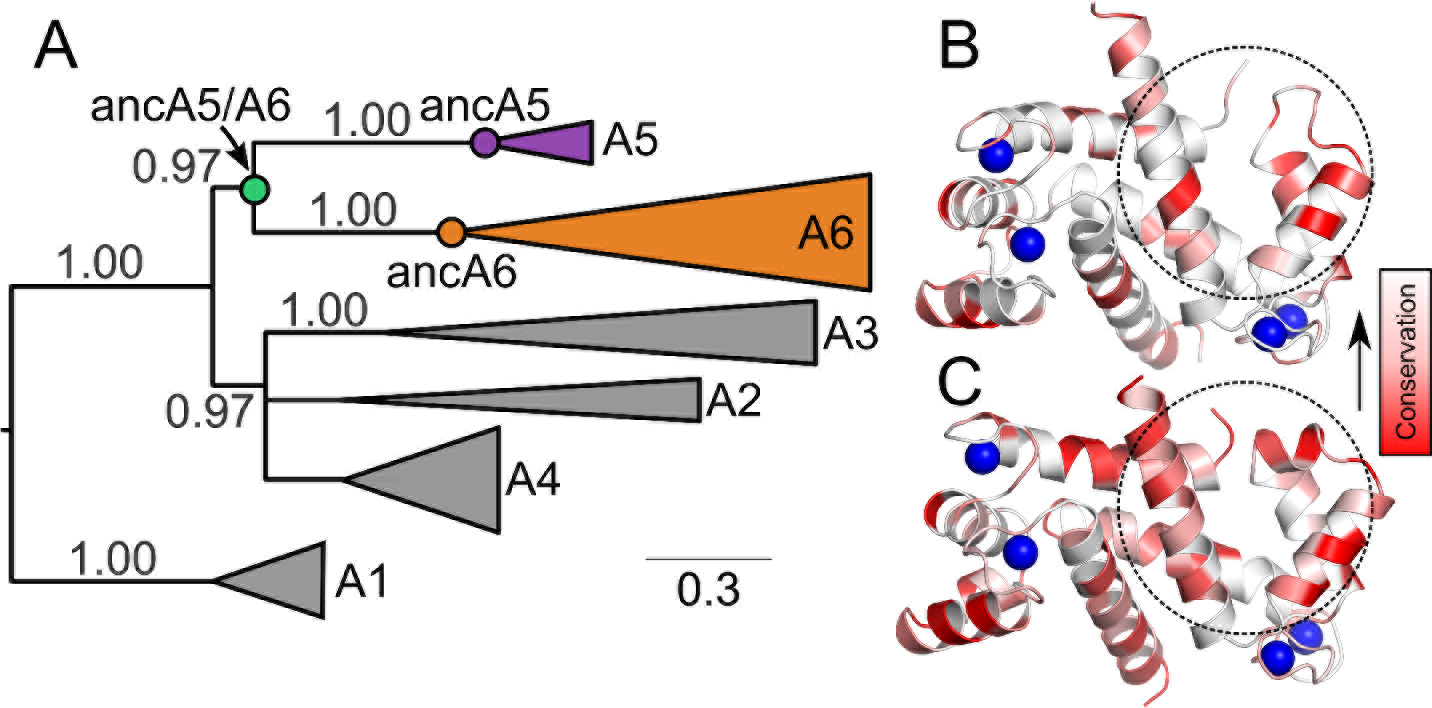
S100A5 and S100A6 arose by gene duplication in the amniote ancestor. A) Maximum likelihood phylogeny for S100A5, S100A6 and their close homologs. Wedges denote collections of paralogs (S100A1, S100A2, S100A3, S100A4, S100A5, or S100A6). Wedge height corresponds to the number of sequences and wedge length to the longest branch in that clade. SH supports, estimated using an approximate likelihood ratio test (76), are shown above the branches. Scale bar shows branch length in substitutions per site. Reconstructed ancestors are denoted with circles. All proteins, with the exception of those in the A1 clade, are taken from amniotes. A1 contains S100 proteins from bony vertebrates and was used as an out-group to root the tree. Panels B and C show relative conservation of residues across amniote paralogs mapped onto the structures of hA5 (2KAY, (46)) and hA6 (1K96, (77)). Colors denote conservation from < 20 % (dark red) to 100 % white. Sequences were taken from the alignment used to generate the phylogeny in panel A. Dashed circles denote the peptide binding surface for one of the two chains. Blue spheres show the location of bound *Ca*^2+^ in the structures.

To test the prediction that specificity has fluctuated over time, we expressed and purified S100A5 and S100A6 orthologs from human, mouse (*Mus musculus*), tasmanian devil (*Sarcophilus harrisii*), American alligator (*Alligator mississippiensis*), and chicken (*Gallus gallus*). We then characterized the peptide binding specificity of these S100A5 and S100A6 orthologs against four peptides: A5cons, A6cons, SIP, and NCX1 (Fig 4A). We selected these peptides because there is direct evidence that these peptides bind at the canonical binding interface (Fig 2, as well as (41, 42)). Surprisingly, we found that the S100A5 and S100A6 clades exhibited broadly similar, ortholog-specific binding specificity (Fig 4A). All S100A5 orthologs bound NCX1, A5cons, and A6cons, but not SIP. In contrast, all S100A6 orthologs bound SIP and A6cons, but not A5cons. The only labile character is NCX1 binding to S100A6. The sauropsid and marsupial S100A6 orthologs bound NCX1, but not the eutherian mammal representatives. We also characterized binding of these peptides to human S100A4 as an outgroup. Binding for this protein was intermediate between the S100A5 and S100A6 clades: it bound A5cons and A6cons, but not SIP or NCX1. Thermodynamic parameters for these binding experiments are given in Table S2-S5. Representative ITC traces for each protein are shown in Fig S2.

**Fig 4.**
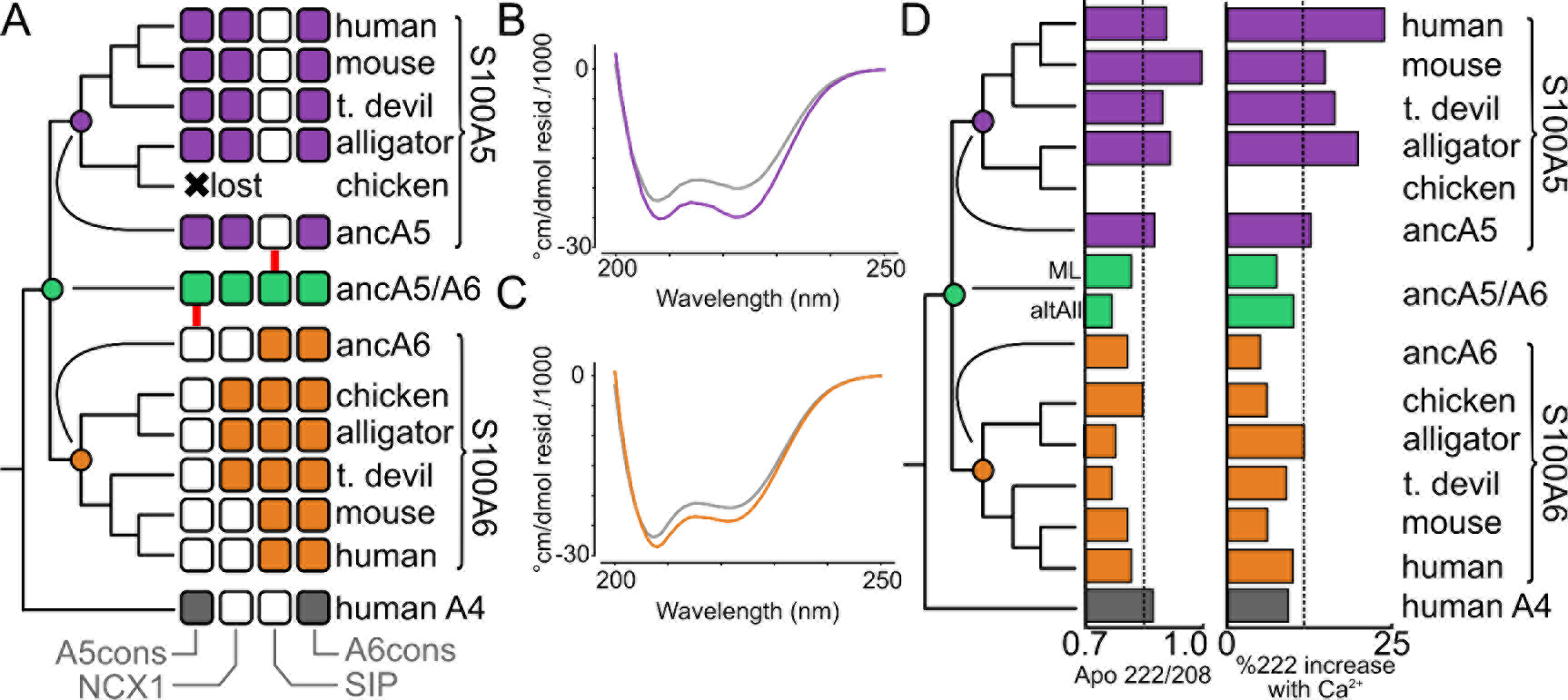
S100A5 and S100A6 paralogs exhibit conserved properties. A) Peptide binding specificity mapped onto the phylogenetic tree as a collection of binary characters. Each square denotes binding of a specific peptide to an ortholog sampled from the species indicated at right. Squares are filled if binding was observed by ITC. Ancestors are shown in the middle, with red arrows indicating changes that occurred after duplication that were then conserved across orthologs. The results for ancA5/A6 were identical for both the ML and “altAll” ancestors. Full thermodynamic parameters are in Table S2-S5. B) Far-UV spectra for apo (gray) and *Ca*^2+^-bound (purple) hA5. C) Far-UV spectra for apo (gray) and *Ca*^2+^-bound (orange) hA6. D) Spectroscopic properties mapped onto the phylogeny. The left column shows the ratio of absorbance at 222 nm/208 nm for the apo protein. The right column shows the percentage increase in signal at 222 nm upon addition of *Ca*^2+^. Dashed lines show the mean values across all experiments. Raw spectra are given in Fig S3.

The strong conservation of peptide binding suggested that other features—such as structural features—might be conserved between paralogs as well. To test for this, we characterized the secondary structure and response to *Ca*^2+^ for all proteins using far-UV circular dichroism (CD) spectroscopy. A *Ca*^2+^-driven change in a-helical secondary structure is a conserved feature of S100 proteins (26, 37). We asked whether this behavior was conserved across orthologs, which would indicate similar structural properties. As with peptide binding, we found that the CD spectrum and response to *Ca*^2+^ were diagnostic within each clade (Fig 4B-D, Fig S3). S100A5 orthologs exhibited deep minima at 208 and 222 nm, corresponding to a largely α-helical secondary structure (Fig 4B,D). This signal increased upon addition of saturating *Ca*^2+^, consistent with the ordering of the C-terminus of the human protein reported by NMR (46). In contrast, all S100A6 orthologs exhibited a deeper minimum at 208 nm, likely corresponding to a mixture of *α*-helical and random coil secondary structure. The secondary structure of these proteins changed comparatively little on addition of *Ca*^2+^(Fig 4C,D).

### Specificity evolved from an apparently promiscuous ancestor

Surprisingly, despite the diversity of peptides that bind to each paralog, peptide binding specificity is conserved across across paralogs. We next asked whether these proteins exhibited comparable evolutionary patterns to those observed in high-specificity proteins, such as the partitioning of ancestral binding partners along duplicate lineages (20–22). Using our phylogeny, we used ancestral sequence reconstruction (ASR) to reconstruct the last common ancestors of S100A5 orthologs (ancA5) and S100A6 orthologs (ancA6) (49). These proteins were well reconstructed, having mean posterior probabilities of 0.93 and 0.96, respectively. Their sequences are given in File S2. We expressed and purified both of these proteins. We found that they shared similar secondary structures and *Ca*^2+^-binding responses with their descendants by far-UV CD (Fig 4C). We then measured binding to the suite of four peptides described above using ITC. These ancestors gave the pattern we would expect given the binding specificities of the derived proteins (Fig 4D). AncA5 is indistinguishable from a modern S100A5 ortholog, binding A5cons, A6cons, and NCX1, but not SIP (Fig 4D). AncA6 also behaves as expected, binding A6cons and SIP, but not A5cons. It does not bind NCX1, consistent with this character being labile in the S100A6 lineage (Fig 4D).

We next characterized the last common ancestor S100A5 and S100A6 (ancA5/A6). This reconstruction had a mean posterior probability of 0.83 (File S2). AncA5/A6 has a secondary structure content identical to ancA6 and the S100A6 descendants. It also responds to *Ca*^2+^ in a similar fashion (Fig 4C, Fig S2). Unlike any modern protein, however, ancA5/A6 binds to all four peptides (Fig 5). To verify that this result was not an artifact of the reconstruction, we also made an “AltAll” ancestor of ancA5/A6 in which we swapped all ambiguous sites in the maximum-likelihood ancestor with their next most likely alternative (50) (File S2, methods). This protein is quite different than ancA5/A6—differing at 21 of 93 sites—but the binding profile for the four peptides was identical to the maximum-likelihood ancestor. Thermodynamic parameters for these binding experiments are given in Table S2-S5.

**Fig 5.**
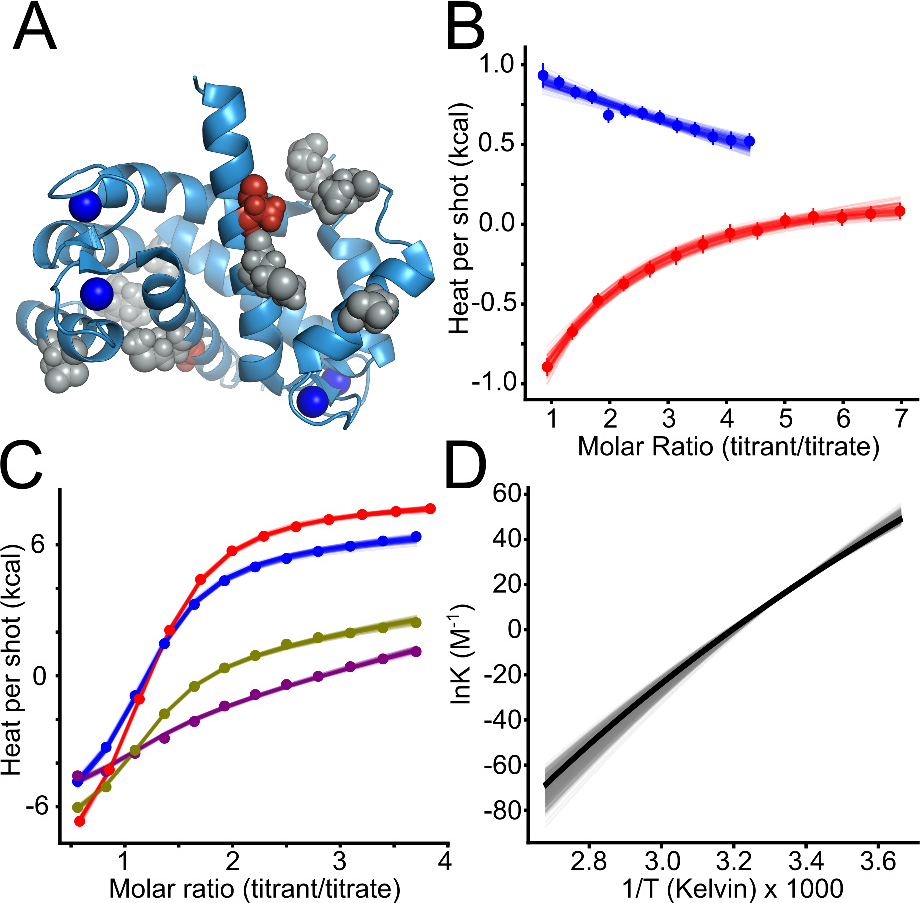
Small changes are sufficient to alter binding specificity at the interface. A) *Ca*^2+^-bound structure of human S100A5 (2KAY) (46) with ancestral reversions marked in gray (no effect on SIP binding) and red (A83, which causes SIP binding). Blue spheres are *Ca*^2+^ ions. B) ITC traces showing titration of SIP onto hA5 A83m (red) versus wildtype hA5 (blue). Points are integrated heats extracted from each shot. Lines are 100 different fit solutions drawn from the fit posterior probability distributions. For the hA5/A83m curve, we used a single-site binding model. For the hA5 curve, we used a blank dilution model, where the linear slope is indicative of peptide dilution without binding. C) ITC traces for titrations of A5cons onto hA5 for as a function of temperature: 10°*C* (purple), 15°*C* (green), 20°*C* (blue), and 25°*C* (red). Points are integrated heats extracted from each shot. Lines are 100 different fit solutions drawn from the fit posterior probability distributions for a global Van’t Hoff model optimized on all four experiments simultaneously. D) Van’t Hoff plot showing temperature dependence of *1n*(*K*) determined from global fit in panel C. Thick black line shows Maximum Likelihood curve, gray lines are 500 curves drawn from the posterior distribution of the Bayesian fit.

### Binding specificity can be changed with a single mutation

Our work revealed that S100A5 and S100A6, despite having low overall specificity, display the same basic evolutionary patterns as high-specificity proteins (20, 22, 23): they exhibit conserved partners across modern orthologs and display a pattern of subfunctionalization from a less specific ancestor. While suggestive, this does not establish that there are functional constraints on specificity. Another possibility is that switching specificity is intrinsically difficult, and that the pattern we observe reflects this difficulty rather than selective pressure to maintain a particular specificity profile.

To distinguish these possibilities, we attempted to shift the binding specificity of hA5 by introducing mutations at the binding interface. We selected five historical substitutions that occurred along the branch between ancA5/A6 and ancA5: e2A, i44L, k54D, a78M, m83A (with the ancestral amino acid in lowercase and modern amino acid in uppercase). We chose these substitutions using three criteria: 1) the ancestral amino acid was conserved in S100A6 orthologs, 2) the derived amino acid was conserved in S100A5 orthologs, 3) and the mutations were located at the peptide binding interface. Fig 5A shows the positions of candidate substitutions mapped onto the structure of hA5 (46).

We reversed each of these sites individually to the ancestral state in hA5. We then measured binding of two clade-specific peptides, SIP and A5cons, to each mutant using ITC (Table S6). We found that reverting a single substitution (A83m) to its ancestral state in hA5 enabled it to bind the SIP peptide (Fig 5B). This reversion does not compromise binding to A5cons, thus recapitulating the ancestral specificity (Table S3). Reversion to the ancestral methionine at residue 83 likely makes more favorable hydrophobic packing interactions with the SIP peptide than the extant alanine. This demonstrates that a single mutation at the peptide binding interface is capable of shifting specificity in S100A5. None of the remaining four ancestral reversions led to measurable changes in A5cons or SIP binding. Amino acids at these positions either do not interact with these peptides, or the ancestral and derived amino acids interact in roughly equivalent fashion.

Another way to view specificity is in terms of binding mechanism. If binding affinity is mostly due to the hydrophobic effect, we would predict it would be relatively easy to alter binding by small changes to packing interactions. To test for relative contributions of the hydrophobic effect versus polar contacts to binding affinity, we did a van’t Hoff analysis for the binding of A5cons to hA5. We performed ITC at temperatures ranging from 10 °*C* to 25 °*C* and then globally fit van’t Hoff models to the binding isotherms (Fig 5C-D). We first attempted fits using a fixed enthalpy of binding 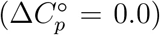, but the fits did not converge. When we allowed 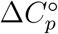 to float, we found it was negative (−0.40 ≤ −0.36 ≤ −0.32 *kcal* · *mol*^−1^ · *K*^−1^), indicating that binding is driven by the hydrophobic effect (51). This observation is consistent with binding at the hydrophobic surface exposed by the *Ca*^2+^-induced conformational change (46) and may help to explain why specificity can be readily altered via a single substitution in the interface.

## Discussion

Our work highlights the paradoxical nature of peptide binding specificity for these low-specificity S100 proteins. The binding interface has low specificity, interacting with very diverse peptides with no obvious binding motif (Fig 1B). Further, the specificity is fragile, and can be altered with a single point mutation (Fig 5). One might therefore conclude that this binding specificity is only weakly constrained. In contrast, binding specificity has been conserved over 320 million years along both lineages, exhibiting a pattern of subfunctionalization similar to what has been observed previously for the evolution of high-specificity proteins (Fig 4). This strongly points to the binding specificity being important, despite being very broad.

### Low specificity through a hydrophobic interface

The binding specificity of these proteins is likely driven almost entirely by shape complementarity and packing. The protein interface exposed on *Ca*^2+^ binding is hydrophobic and likely makes few protein-peptide polar contacts. This prediction is validated, at least for the hA5/A5cons interaction, by the negative 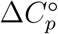 on binding, pointing to an important contribution from the hydrophobic effect on binding (Fig 5C). The lack of polar contacts is the likely explanation for the low specificity of the interface. Peptides need only match hydrophobicity and packing, meaning that a large number of possible peptides bind with similar affinity.

The hydrophobic nature of the interface explains the low specificity, but makes the conservation of specificity over 320 million years quite surprising. There is likely no diagnostic set of polar contacts that can be conserved maintain specificity. It should therefore be straightforward to change specificity with minimal perturbation. Indeed, we found that a single mutation, from a small to a large hydrophobic amino acid, is able to switch the specificity of the interface (Fig 5A). Yet, over evolutionary time, binding specificity—at least for this set of targets—has been maintained (Fig 4). Amazingly, this is achieved without strict conservation of the binding site. The peptide binding region does not exhibit higher conservation than other residues in either S100A5 or S100A6 (Fig 3B-C).

Our work shows that protein binding specificity is likely an important feature of these proteins, but does not reveal the set of biological targets for S100A5 and S100A6. Identifying these targets will require further experiments. This could include coupling S100A5 and S100A6 knockouts to proteomics or transcriptomics, pull downs followed by proteomics, and/or large-scale screens of peptide targets via a technique like phage display. We also anticipate that external factors—such as coexpression, large complex assembly, and subcellular localization—will add critical additional layers of specificity to the low-specificity binding interfaces of these proteins. Understanding the interplay between the biochemical specificity and these external factors will be important for dissecting the biology of these proteins.

### S100s may allow the evolution of new calcium regulation

The existence of a conserved set of binding partners also has intriguing implications for the evolution of *Ca*^2+^ signaling pathways in vertebrates. This can be seen by contrasting S100 proteins with calmodulin, a protein that also exposes a protein interaction surface and regulates the activity of target proteins in response to *Ca*^2+^ (2). It has been proposed that calmodulin provides a universal *Ca*^2+^ response across tissues, while S100 proteins allow for fine-tuned, tissue-specific responses (26, 27). Our results allow us to extend this idea along an evolutionary axis.

Our results suggest that S100 proteins may provide a minimally pleiotropic pathway for the evolution of new *Ca*^2+^ regulation. Calmodulin is broadly expressed across tissues. As a result, a mutation that causes a protein to interact with calmodulin will have the same effect in all tissues where that protein is expressed. This could lead to unfavorable pleiotropic effects that prevent fixation of the mutation. In contrast, S100 proteins have highly differentiated tissue expression. S100A5, for example, is expressed almost exclusively in olfactory tissues. This means that a protein that acquires an interaction with S100A5 will do so only in olfactory tissue, with minimal pleiotropic effects in other tissues. The pattern of subfunctionalization we observed is consistent with this idea (Fig 4D), as subfunctionalization is one way to escape adaptive conflict that arises due to pleiotropic effects of mutations (52, 53). This is only possible because S100A5 evolved a distinct binding profile relative to S100A6 (and presumably other S100 proteins), meaning that acquisition of a new S100A5 interaction does not imply an interaction with a large number of other S100 proteins, which would itself lead to extensive pleiotropy.

Additionally, our results suggest that S100 proteins would provide a much simpler path for the evolution of new *Ca*^2+^ regulation than calmodulin. The calmodulin sequence has been conserved for over a billion years and is basically unchanged across fungi and animals. As a result, evolution of a new calmodulin-regulated target requires that the target change its sequence to bind to calmodulin. This would likely mean that slowly evolving proteins would not be able to evolve *Ca*^2+^ regulation, as neither the calmodulin nor possible new target would be able to acquire the necessary mutations to form the new interaction. In contrast, S100 proteins are evolving rapidly. For example, human S100A5 and S100A6 only exhibit 53% sequence identity, despite sharing an ancestor ≈ 320 million years ago. This means that, particularly after gene duplication, S100 proteins can acquire new interactions through mutations to the S100 itself. This would allow them to capture slowly evolving target proteins, opening a different avenue for the evolution of *Ca*^2+^ regulation that would not be accessible by calmodulin alone.

### Evolution of low-specificity proteins

Our results also shed light on the evolution of low specificity proteins in general. Many proteins besides S100 proteins exhibit low specificity including other signaling proteins (2, 12), hub proteins (3, 6, 9, 11), and many others (1, 4, 5, 8, 10). Further experiments will be required to determine the generality of our observations for low-specificity proteins, but our work suggests that low-specificity proteins can evolve with similar dynamics to the high-specificity proteins that have been studied in detail. Partners for low-specificity proteins can be strongly conserved and evolve by subfunctionalization, just like a high-specificity protein.

One important question is whether S100A5 and S100A6 did, indeed, gain specificity over time. The current study, like many others (17, 20, 54–58), revealed an ancestral protein that appears less specific than its descendants. Some have proposed this is a general evolutionary trend (17, 54, 58). Caution is warranted before interpreting these data as evidence for this hypothesis. We selected a small set of peptides to study; therefore, other patterns may be consistent with our observations. For example, it could be that the proteins both acquired more peptides that we did not sample in this experiment (actual neofunctionalization), while becoming more specific for the chosen set of targets (apparent subfunctionalization). Particularly given the large number of targets for these proteins, distinguishing these possibilities will require an unbiased, high-throughout approach to measuring specificity. Advances in high-throughput protein characterization have made such experiments tractable (59–63). With the right method, we will be able to resolve whether the shifts in specificity we observed indeed reflect increased specificity over evolutionary time, or instead the small size of the binding set we investigated.

Whatever the precise evolutionary process, our results reveal that S100 proteins—despite binding diverse peptides at a low-specificity hydrophobic interface—have maintained the same binding profile for the last 320 million years. Low-specificity does not imply no specificity, nor a lack of evolutionary constraint.

## Acknowledgements

We would like to thank members of the Harms group for useful discussions. We would also like to thank Patrick Reardon of the OSU NMR core facility for his assistance with using the 800 MHz instrument. This work was funded by NIH R01GM117140 (MJH) and NIH 7T32GM007759 (LCW). The Oregon State University NMR Facility is funded in part by the National Institutes of Health, HEI Grant 1S10OD018518 and by the M. J. Murdock Charitable Trust grant #2014162. The funders had no role in study design, data collection and analysis, decision to publish, or preparation of the manuscript.

## Materials and Methods

### Molecular cloning, expression and purification of proteins

Synthetic genes encoding the S100 proteins and codon-optimized for expression in *E. coli* were ordered from Genscript. The accession numbers for the modern sequences are: *Homo sapiens* S100A5: P33763, S100A6: P06703; *Mus musculus* S100A5: P63084, S100A6: P14069; *Sarcophilus harrisii* S100A5: G3W581, S100A6: G3W4S8; *Alligator mississippiensis* S100A5: XP_006264408.1, S100A6: XP_006264409.1; *Gallus gallus* S100A6: Q98953. All accession numbers are for the uniprot database (64), with the exception of the *Alligator mississippiensis* accessions, which are for the NCBI database (65).

Genes were sub-cloned into a pET28/30 vector containing an N-terminal His tag with a TEV protease cleavage site (Millipore). Expression was carried out in Rosetta (DE3) pLysS *E. coli* cells. 1.5 L cultures were inoculated at a 1:100 ratio with saturated overnight culture. E.coli were grown to high log-phase (*OD*_600_≈0.8-1.0) with 250rpm shaking at 37°*C*. Cultures were induced by addition of 1 mM IPTG along with 0.2% glucose overnight at 16°C. Cultures were centrifuged and the cell pellets were frozen at −20°*C* and stored for up to 2 months. Lysis of the cells was carried out via sonication in 25mM Tris, 100mM NaCl, 25mM imidazole, pH 7.4.

Purification of all S100s used in this study was carried out as follows. The initial purification step was performed using a 5 mL HiTrap Ni-affinity column (GE Health Science) on an Äkta PrimePlus FPLC (GE Health Science). Proteins were eluted using a 25mL gradient from 25-500mM imidazole in a background buffer of 25mM Tris, 100mM NaCl, pH 7.4. Peak fractions were pooled and incubated overnight at 4°*C* with ≈1:5 TEV protease (produced in the lab). TEV protease removes the N-terminal His-tag from the protein and leaves a small Ser-Asn sequence N-terminal to the wildtype starting methionine. Next hydrophobic interaction chromatography (HIC) was used to purify the S100s from remaining bacterial proteins and the added TEV protease. Proteins were passed over a 5 mL HiTrap phenyl-sepharose column (GE Health Science). Due to the Ca^2+^-dependent exposure of a hydrophobic binding, the S100 proteins proteins adhere to the column only in the presence of Ca^2+^. Proteins were pre-saturated with 2mM *Ca*^2+^ before loading on the column and eluted with a 30mL gradient from 0mM to 5mM EDTA in 25mM Tris, 100mM NaCl, pH 7.4. Peak fractions were pooled and dialyzed against 4 L of 25 mM Tris, 100 mM NaCl, pH 7.4 buffer overnight at 4°*C* to remove excess EDTA. The proteins were then passed once more over the 5 mL HiTrap Ni-affinity column (GE Health Science) to removed any uncleaved His-tagged protein. The cleaved protein was collected in the flow-through. Finally, protein purity was examined by SDS-PAGE. If any trace contaminants appeared to be present we performed anion chromatography with a 5mL HiTrap DEAE column (GE). Proteins were eluted with a 50mL gradient from 0-500mM NaCl in 25mM Tris, pH 7.0-8.5 (dependent on protein isolectric point) buffer. Pure proteins were dialyzed overnight against 2L of 25mM TES (or Tris), 100mM NaCl, pH 7.4, containing 2 g Chelex-100 resin (BioRad) to remove divalent metals. After final purification step, the purity of proteins products was assessed by SDS PAGE and MALDI-TOF mass spectrometry to be > 95. Final protein products were flash frozen, dropwise, in liquid nitrogen to form frozen spherical pellets and stored at −80°*C*. Protein yields were typically on the order of 25mg/1.5L of culture.

### Isothermal titration calorimetry

ITC experiments were performed in 25 mM TES, 100mM NaCl, 2mM CaCl2, 1mM TCEP, pH 7.4. Although most experiments were performed at 25°*C*, some were done at cooler temperatures depending to ensure measurable binding heats and sufficient curvature for fitting. Samples were equilibrated and degassed by centrifugation at 18, 000*xg* at the experimental temperature for 30 minutes. Peptides (GenScript, Inc.) were dissolved directly into the experimental buffer prior to each experiment. All experiments were performed at on a MicroCal ITC-200 or a MicroCal VP-ITC (Malvern). Gain settings were determined on a case-by-case basis to ensured quality data. A 750 rpm syringe stir speed was used for all ITC-200 experiments while 400rpm speed was used for experiments on the VP-ITC. Spacing between injections ranged from 300s-900s depending on gain settings and relaxation time of the binding process. These setting were optimized for each binding interaction that was measured. Titration data were fit to a single-site binding model using the Bayesian fitter in pytc. For each protein/peptide combination, one clean ITC trace was used to fit the binding model. Negative results were double-checked to ensure accuracy. Some were done at lower temperatures (10°*C* or 15°*C*) to confirm lack of binding, because peptide binding enthalpy should be dependent on temperature.

### 2D HSQC NMR experiments

We collected 2D ^1^*H* –^15^ *N* TROSY-HSQC NMR spectra for 2 *mM* hA5 in the presence of *Ca*^2+^ alone and with the addition of the 2 *mM* A5cons. We also collected the spectra of 0.5 *mM* hA5 with the addition of 0.5 *mM* A6cons peptide, which was done at lower concentration due to poorer solubility of A6cons in the aqueous buffer. We transfered published assignments to the *Ca*^2+^-alone spectrum (BMRB: 16033, (46)), and then used 3D NOESY-TROSY spectra to verify the assignments. We were able to unambiguously assign 76 peaks of the 91 non-proline amino acids in the *Ca*^2+^-bound form. We then added saturating A5cons or A6cons peptide to the sample and remeasured the TROSY-HSCQ spectrum. We then noted which peaks had either shifted or entered intermediate exchange upon addition of the peptide. Of the 76 unambiguously assigned non-proline amino acids 26 shifted or disappeared in the A5cons-bound form, and 35 shifted or disappeared in the A6cons bound form.

All NMR experiments were performed at 25 °*C* on an 800 MHz (18.8T) Bruker spectrometer at Oregon State University. TROSY spectra were collected with 32 transients, 1024 direct points with a signal width of 12820, and 256 indirect points with a signal width of 2837 Hz in ^15^*N*. NOESY-TROSYs were run with 8 transients, non-uniform sampling with 15% of data points used, and a 150 ms mixing time. All spectra were processed using NMRPipe (66); data were visualized and assignments transferred using the CCPNMR analysis program (67).

### Far-UV CD spectroscopy

Far-UV circular dichroism spectra (200-250nm) were collected on a J-815 CD spectrometer (Jasco) with a 1 mm quartz cell (Starna Cells, Inc.). We prepared 20-40 *μM* samples in a Chelex (Bio-Rad) treated, 25mM TES (Sigma), 100mM NaCl (Thermo Scientific) buffer at pH 7.4. Samples were centrifuged at 18,000 x g at 25°*C* in a temperature-controlled centrifuge (Eppendorf) before experiments. Spectra were measured in the absence and presence of saturating Ca^2+^. Reversibility of Ca^2+^-induced structural changes was confirmed by subsequently adding a molar excess of EDTA to the Ca^2+^-saturated samples and repeating the measurements. Five scans were collected for each condition and averaged to minimize noise. A buffer blank spectrum was subtracted with the built-in subtraction feature in the Jasco spectra analysis software. Raw ellipticity was later converted into mean molar ellipticity based on the concentration and residue length of each protein. These calculations were performed on the buffer-blanked data.

### Preparation of biotinylated proteins for phage display

A small amount of the purified proteins were biotinylated in the following way using the EZ-link BMCC-biotin system (ThermoFisher Scientific). This kit used a maleimide linker to attach biotin at a Cys residue on the protein. ≈1mg BMCC-biotin was dissolved directly in 100% DMSO to a concentration of 8mM for labeling. Proteins were exchanged into 25mM phosphate, 100mM NaCl, pH 7.4 using a Nap-25 desalting column (GE Health Science) and degassed for 30 minutes at 25°*C* using a vacuum pump (Malvern Instruments). While stirring at room temperature, 8mM BMCC-biotin was added dropwise to a final 10X molar excess. Reaction tubes were sealed with PARAFILM (Bemis) and the maleimide-thiol reactions were allowed to proceed for 1 hour at room temperature with stirring. The reactions were then transferred to 4°*C* and incubated with stirring overnight to allow completion of the reaction. Excess BMCC-biotin was removed from the labeled proteins by exchanging again over a Nap-25 column (GE Health Science), and subsequently a series of 3 concentration-wash steps on a NanoSep 3K spin column (Pall corporation), into the Ca-TeBST loading loading buffer. Complete labeling was confirmed by MALDI-TOF mass spectrometry by observing the ≈540Da shift in the protein peak. Final stocks of labeled proteins were prepared at 10 *μM* by dilution into the loading buffer.

### Phage display panning

Phage display experiments were performed using the PhD-12 peptide phage display kit (NEB). All steps involving the pipetting of phage-containing samples was done using filter tips to prevent cross-contamination (Rainin). 100μL samples containing phage (2.5*x*10^10^ *PFU*) and biotin-protein 0.01 *μM* (or 0.01 *μM* biotin in the negative control) and 50 *μM* peptide competitor (in competitor samples) were prepared at room temperature in a background of Ca-TeBST loading buffer (25mM TES, 100mM NaCl, 2mM *CaCl*_2_, 0.01% Tween-20, pH 7.4) to ensure saturation of the S100s with Ca^2+^. Samples were incubated at room temperature for 1hr. Each sample was then applied to one well of a 96-well high-capacity streptavidin plate (previously blocked using PhD-12 kit blocking buffer and washed 6X with 150 *μL* loading buffer). Samples were incubated on the plate with gentle shaking for 20min. 1 *μL* of 10mM biotin (NEB) was then added to each sample on the plate and incubated for an additional five minutes to compete away purely biotin-dependent interactions. Samples were then pulled from the plate carefully by pipetting and discarded. Each well was washed 5X with 200 *μL* of loading buffer by applying the solution to the well and then immediately pulling off by pipetting. Finally, 100 *μL* of EDTA-TeBST (25mM TES, 100mM NaCl, 5mM EDTA, 0.01% Tween-20, pH 7.4) elution buffer was applied to each well and the plate was incubated with gentle shaking for 1hr at room temperature to elute. Two replicates of the experiment were performed with each protein.

Eluates were pulled from the plate carefully by pipetting and stored at 4°*C* Eluates were titered to quantify enrichment as follows. Serial dilutions of the eluates from 1 : 10-1 : 10^6^were prepared in LB medium. These were used to inoculate 200 *μL* aliquots of mid-log-phase ER2738 *E. coli* (NEB) by adding 10 *μL* to each. Each 200 *μL* aliquot was then mixed with 3mL of pre-melted top agar, applied to a LB/agar/XGAL/IPTG (Rx Biosciences) plate, and allowed to cool. The plates were incubated overnight at 37°*C* to allow formation of plaques. The next morning, blue plaques were counted and used to calculate PFU/mL phage concentration. Enrichment was calculated as a ratio of experimental samples to the biotin-only negative control.

For subsequent rounds of panning the eluates were amplified as follows. 20mL 1:100 dilutions of an ER2738 overnight culture were prepared. Each 20mL culture was inoculated with one entire sample of remaining phage eluate. The cultures were incubated at 37°*C* with shaking for 4.5 hours to allow phage growth. Bacteria were then removed by centrifugation and the top 80% of the culture was removed carefully with a filtered serological pipette and transferred to a fresh tube containing 1/6 volume of PEG/NaCl (20% w/v PEG-8000, 2.5M NaCl). Samples were incubated overnight at 4°*C* to precipitate phage. Precipitated phage were isolated by centrifugation and subsequently purified by an additional PEG/NaCl precipitation on ice for 1hr. Isolated phage were resuspended in 200 *μL* each sterile loading buffer, titered to measure PFU/mL, and stored at 4°*C* for use in the next panning round. This process was repeated for 3 total rounds of panning. Plaques were pulled from final reound eluate titer plates and amplified in 1mL ER2738 culture for 4.5 hours. ssDNA was isolated from the phage cultures using the Qiagen M13 spin kit. 10 plaques per replicate experiment were Sanger sequenced (GeneWiz, Inc.). These plaque sequences were used to construct the A5cons and A6cons consensus peptides.

### Phylogenetics and ancestral reconstruction

We used targeted BLAST searches to build an database of 49 S100A2-S100A6 sequences sampled from across the amniotes, as well as six telost fish S100A1 sequences as an outgroup. We attempted to achieve even taxonomic sampling across amniotes. Database accession numbers are in Table S7. We used MSAPROBS for the initial alignment (68), followed by manual refinement. Our final alignment is available as a supplemental stockholm file (File S1).

We constructed our phylogenetic tree using the EX/EHO+Г_8_ model, which incorporates information about secondary structure and solvent accessibility into the phylogenetic inference (47). We assigned the secondary structure and solvent accessibility of each site using 115 crystallographic and NMR structures of S100A2, S100A3, S100A4, S100A5 and S100A6 paralogs: 1a03, 1a4p, 1b4c, 1bt6, 1cb1, 1cdn, 1cfp, 1clb, 1cnp, 1ig5, 1igv, 1irj, 1jwd, 1k2h, 1k8u, 1k9p, 1ksm, 1kso, 1m31, 1mq1, 1nsh, 1ozo, 1psb, 1psr, 1sym, 1uwo, 1yur, 1yus, 2bca, 2bcb, 2cnp, 2cxj, 2jpt, 2jtt, 2k8m, 2kax, 2ki4, 2ki6, 2kot, 2l0p, 2l50, 2l5x, 2le9, 2lhl, 2llt, 2llu, 2lnk, 2pru, 2rgi, 2wc8, 2wcb, 2wce, 2wcf, 3ko0, 3nsi, 3nsk, 3nsl, 3nso, 3nxa, 1b1g, 1e8a, 1gqm, 1j55, 1k96, 1k9k, 1mho, 1mr8, 1odb, 1qlk, 1xk4, 1xyd, 1yut, 1yuu, 1zfs, 2egd, 2h2k, 2h61, 2k7o, 2kay, 2l51, 2psr, 2q91, 2wnd, 2wor, 2wos, 2y5i, 3c1v, 3cga, 3cr2, 3cr4, 3cr5, 3czt, 3d0y, 3d10, 3gk1, 3gk2, 3gk4, 3hcm, 3icb, 3iqo, 3lk0, 3lk1, 3lle, 3m0w, 3psr, 3rlz, 4duq, 1mwn, 1qls, 2k2f, 2kbm, 3iqq, 3rm1, 3zwh, 4eto. We calculated the secondary structure for each site using DSSP and the solvent accessibility using NACCESS (69, 70). To remove redundancy—whether from identical sequences solved under slightly different conditions or from the multiple models in the NMR models—we took the majority rule consensus secondary structure and the average solvent accessibility for all structures with identical sequences before doing averages across unique sequences. We then assigned the secondary structure for each column using a majority-rule across unique sequences. We assigned the solvent accessibility as the average across unique sequences at that site. Our structural annotation is available in our alignment stockholm file (File S1).

We then constructed our tree using the EX/EHO+Г_8_ model (47), enforcing correct species relationships within groups of orthologs (71). We compared the final likelihood of this tree to trees generated using LG+Г_8_ and JTT+Г_8_ models (72, 73). Although the EX/EHO model has seven more floating parameters than either LG or JTT, the final tree had a log-likelihood 61 units higher than the next-best model. An AIC test strongly supports the more complex model (*p* = 3 × 10^−30^). One important output from an EX/EHO calculation is χ, a term that measures the fraction of sites that use the structural models relative to a linear combination of all of them (47). For our analysis, χ = 0.72. We rooted the tree using the S100A1 sequences, which included S100s from several bony fishes.

To reconstruct ancestors using the EX/EHO+Г_8_ model, we used PAML to reconstruct ancestors using each of the six possible EX/EHO matrices (49, 74), as well as their linear combination. We then mixed the resulting ancestral posterior probabilities using the secondary structure calls and apparent accessibility at each site, as well as χ (see Equation 3 in (47)). The code implementing this approach is posted on github: https://github.com/harmslab/exexho_phylo_mixer. We assigned gaps using parsimony. We generated the AltAll sequence as described in Eick et al (50). This incorporates uncertainty in the reconstruction by taking the next-best reconstruction at each all ambiguous sites. We took each site at which the posterior probability of the next-best reconstruction was greater than 0.20 and the introduced that alternate reconstruction at the site of interest. Our AltAll sequence differed from the maximum likelihood sequence at 21 positions (24% of sites). File S2 has the posterior probabilities of reconstructions at each site in the ancestor, as well as the final sequences characterized.

## References

[1] Kreegipuu A, Blom N, Brunak S, Järv J (1998) Statistical analysis of protein kinase specificity determinants. FEBS Letters 430(1):45–50.

[2] Chin D, Means AR (2000) Calmodulin: a prototypical calcium sensor. Trends in Cell Biology 10(8):322–328.

[3] Ekman D, Light S, Björklund [U+FFFD], Elofsson A (2006) What properties characterize the hub proteins of the protein-protein interaction network of Saccharomyces cerevisiae? Genome Biology 7(6):R45.

[4] Schreiber G, Keating AE (2011) Protein binding specificity versus promiscuity. Current Opinion in Structural Biology 21(1):50–61.

[5] Nakahara KS, et al. (2012) Tobacco calmodulin-like protein provides secondary defense by binding to and directing degradation of virus RNA silencing suppressors. Proceedings of the National Academy of Sciences 109(25):10113–10118.

[6] Bertolazzi P, Bock ME, Guerra C (2013) On the functional and structural characterization of hubs in protein-protein interaction networks. Biotechnology Advances 31(2):274–286.

[7] Nakagawa S, Gisselbrecht SS, Rogers JM, Hartl DL, Bulyk ML (2013) DNA-binding specificity changes in the evolution of forkhead transcription factors. Proceedings of the National Academy of Sciences 110(30):12349–12354.

[8] Mitchell PS, Emerman M, Malik HS (2013) An evolutionary perspective on the broad antiviral specificity of MxA. Current Opinion in Microbiology 16(4):493–499.

[9] Peleg O, Choi JM, Shakhnovich E (2014) Evolution of Specificity in Protein-Protein Interactions. Biophysical Journal 107(7):1686–1696.

[10] Howard CJ, et al. (2014) Ancestral resurrection reveals evolutionary mechanisms of kinase plasticity. eLife 3:e04126.

[11] Uchikoga N, Matsuzaki Y, Ohue M, Akiyama Y (2016) Specificity of broad protein interaction surfaces for proteins with multiple binding partners. Biophysics and Physicobiology 13:105–115.

[12] Bhattacharya S, Bunick CG, Chazin WJ (2004) Target selectivity in EF-hand calcium binding proteins. Biochimica et Biophysica Acta (BBA) - Molecular Cell Research 1742(1-3):69–79.

[13] Chazin WJ (2011) Relating Form and Function of EF-hand Calcium Binding Proteins. Accounts of chemical research 44(3):171–179.

[14] Harms MJ, Thornton JW (2013) Evolutionary biochemistry: revealing the historical and physical causes of protein properties. Nature reviews. Genetics 14(8):559–571.

[15] McKeown A, et al. (2014) Evolution of DNA Specificity in a Transcription Factor Family Produced a New Gene Regulatory Module. Cell 159(1):58–68.

[16] Boucher JI, Jacobowitz JR, Beckett BC, Classen S, Theobald DL (2014) An atomic-resolution view of neofunctionalization in the evolution of apicomplexan lactate dehydrogenases. eLife 3:e02304.

[17] Khersonsky O, Tawfik DS (2010) Enzyme Promiscuity: A Mechanistic and Evolutionary Perspective. Annual Review of Biochemistry 79(1):471–505.

[18] Copley SD (2012) Toward a Systems Biology Perspective on Enzyme Evolution. The Journal of Biological Chemistry 287(1):3–10.

[19] Wheeler LC, Lim SA, Marqusee S, Harms MJ (2016) The thermostability and specificity of ancient proteins. Current Opinion in Structural Biology 38:37–43.

[20] Eick GN, Colucci JK, Harms MJ, Ortlund EA, Thornton JW (2012) Evolution of Minimal Specificity and Promiscuity in Steroid Hormone Receptors. PLoS Genetics 8(11).

[21] Hudson WH, et al. (2015) Distal substitutions drive divergent DNA specificity among paralogous transcription factors through subdivision of conformational space. Proceedings of the National Academy of Sciences p. 201518960.

[22] Clifton B, Jackson C (2016) Ancestral Protein Reconstruction Yields Insights into Adaptive Evolution of Binding Specificity in Solute-Binding Proteins. Cell Chemical Biology 23(2):236–245.

[23] Alhindi T, et al. (2017) Protein interaction evolution from promiscuity to specificity with reduced flexibility in an increasingly complex network. Scientific Reports 7.

[24] Aakre CD, et al. (2015) Evolving New Protein-Protein Interaction Specificity through Promiscuous Intermediates. Cell 163(3):594–606.

[25] Sayou C, et al. (2014) A Promiscuous Intermediate Underlies the Evolution of LEAFY DNA Binding Specificity. Science 343(6171):645–648.

[26] Marenholz I, Heizmann CW, Fritz G (2004) S100 proteins in mouse and man: from evolution to function and pathology (including an update of the nomenclature). Biochemical and Biophysical Research Communications 322(4):1111–1122.

[27] Donato R, et al. (2013) Functions of S100 Proteins. Current molecular medicine 13(1):24–57.

[28] Sivaraja V, et al. (2006) Copper Binding Affinity of S100a13, a Key Component of the FGF-1 Nonclassical Copper-Dependent Release Complex. Biophysical Journal 91(5):1832–1843.

[29] Carreira CM, et al. (1998) S100a13 Is Involved in the Regulation of Fibroblast Growth Factor-1 and p40 Synaptotagmin-1 Release in Vitro. Journal of Biological Chemistry 273(35):22224–22231.

[30] Yang Z, et al. (2007) S100a12 provokes mast cell activation: a potential amplification pathway in asthma and innate immunity. The Journal of Allergy and Clinical Immunology 119(1):106–114.

[31] Leclerc E, Fritz G, Vetter SW, Heizmann CW (2009) Binding of S100 Proteins to RAGE: An Update. Biochimica et Biophysica Acta (BBA) - Molecular Cell Research 1793(6):993–1007.

[32] Cho CC, Chou RH, Yu C (2016) Pentamidine blocks the interaction between mutant S100a5 and RAGE V domain and inhibits the RAGE signaling pathway. Biochemical and Biophysical Research Communications 477(2):188–194.

[33] Damo SM, et al. (2013) Molecular basis for manganese sequestration by calprotectin and roles in the innate immune response to invading bacterial pathogens. Proceedings of the National Academy of Sciences 110(10):3841–3846.

[34] Santamaria-Kisiel L, Rintala-Dempsey AC, Shaw GS (2006) Calcium-dependent and -independent interactions of the S100 protein family. Biochemical Journal 396(2):201–214.

[35] Streicher WW, Lopez MM, Makhatadze GI (2009) Annexin I and Annexin II N-Terminal Peptides Binding to S100 Protein Family Members: Specificity and Thermodynamic Characterization. Biochemistry 48(12):2788–2798.

[36] Hedges SB, Dudley J, Kumar S (2006) TimeTree: A Public Knowledge-Base of Divergence Times among Organisms. Bioinformatics 22(23):2971–2972.

[37] Wheeler LC, Donor MT, Prell JS, Harms MJ (2016) Multiple Evolutionary Origins of Ubiquitous Cu2+ and Zn2+ Binding in the S100 Protein Family. PloS one 11(10):e0164740.

[38] Leśniak W, S\lomnicki \P, Filipek A (2009) S100a6 - New Facts and Features. Biochemical and Biophysical Research Communications 390(4):1087–1092.

[39] S\lomnicki \P, Nawrot B, Leśniak W (2009) S100a6 Binds P53 and Affects Its Activity. The International Journal of Biochemistry & Cell Biology 41(4):784–790.

[40] van Dieck J, Fernandez-Fernandez MR, Veprintsev DB, Fersht AR (2009) Modulation of the Oligomerization State of P53 by Differential Binding of Proteins of the S100 Family to P53 Monomers and Tetramers. Journal of Biological Chemistry 284(20):13804–13811.

[41] Lee YT, et al. (2008) Structure of the S100a6 Complex with a Fragment from the C-Terminal Domain of Siah-1 Interacting Protein: A Novel Mode for S100 Protein Target Recognition. Biochemistry 47(41):10921–10932.

[42] Liriano MA (2012) Ph.D. (University of Maryland, Baltimore, United States - Maryland).

[43] Knott TK, et al. (2012) Olfactory Discrimination Largely Persists in Mice with Defects in Odorant Receptor Expression and Axon Guidance. Neural development 7(1):17.

[44] McIntyre JC, et al. (2012) Gene Therapy Rescues Cilia Defects and Restores Olfactory Function in a Mammalian Ciliopathy Model. Nature medicine 18(9):1423–1428.

[45] Olender T, et al. (2016) The Human Olfactory Transcriptome. BMC genomics 17(1):619.

[46] Bertini I, et al. (2009) Solution Structure and Dynamics of S100a5 in the Apo and Ca2+-Bound States. JBIC Journal of Biological Inorganic Chemistry 14(7):1097–1107.

[47] Le SQ, Gascuel O (2010) Accounting for Solvent Accessibility and Secondary Structure in Protein Phylogenetics Is Clearly Beneficial. Systematic Biology 59(3):277–287.

[48] Zimmer DB, Eubanks JO, Ramakrishnan D, Criscitiello MF (2013) Evolution of the S100 family of calcium sensor proteins. Cell Calcium 53(3):170–179.

[49] Yang Z, Kumar S, Nei M (1995) A New Method of Inference of Ancestral Nucleotide and Amino Acid Sequences. Genetics 141(4):1641–1650.

[50] Eick GN, Bridgham JT, Anderson DP, Harms MJ, Thornton JW (2017) Robustness of Reconstructed Ancestral Protein Functions to Statistical Uncertainty. Molecular Biology and Evolution 34(2):247–261.

[51] Connelly PR, Thomson JA (1992) Heat Capacity Changes and Hydrophobic Interactions in the Binding of FK506 and Rapamycin to the FK506 Binding Protein. Proceedings of the National Academy of Sciences of the United States of America 89(11):4781–4785.

[52] Des Marais DL, Rausher MD (2008) Escape from adaptive conflict after duplication in an anthocyanin pathway gene. Nature 454(7205):762–765.

[53] Soskine M, Tawfik DS (2010) Mutational effects and the evolution of new protein functions. Nature Reviews Genetics 11(8):572–582.

[54] Zou T, Risso VA, Gavira JA, Sanchez-Ruiz JM, Ozkan SB (2015) Evolution of Conformational Dynamics Determines the Conversion of a Promiscuous Generalist into a Specialist Enzyme. Molecular Biology and Evolution 32(1):132–143.

[55] Carroll SM, Bridgham JT, Thornton JW (2008) Evolution of Hormone Signaling in Elasmobranchs by Exploitation of Promiscuous Receptors. Molecular Biology and Evolution 25(12):2643–2652.

[56] Devamani T, et al. (2016) Catalytic promiscuity of ancestral esterases and hydroxynitrile lyases. Journal of the American Chemical Society 138(3):1046–1056.

[57] Voordeckers K, Pougach K, Verstrepen KJ (2015) How do regulatory networks evolve and expand throughout evolution? Current Opinion in Biotechnology 34(Supplement C):180–188.

[58] Risso VA, Gavira JA, Sanchez-Ruiz JM (2014) Thermostable and promiscuous Precambrian proteins. Environmental Microbiology 16(6):1485–1489.

[59] Carlson CD, et al. (2010) Specificity landscapes of DNA binding molecules elucidate biological function. Proceedings of the National Academy of Sciences 107(10):4544–4549.

[60] Fowler DM, et al. (2010) High-resolution mapping of protein sequence-function relationships. Nature Methods 7(9):741–746.

[61] Ernst A, et al. (2010) Coevolution of PDZ domain-ligand interactions analyzed by high-throughput phage display and deep sequencing. Molecular BioSystems 6(10):1782.

[62] Teyra J, Sidhu SS, Kim PM (2012) Elucidation of the binding preferences of peptide recognition modules: SH3 and PDZ domains. FEBS letters 586(17):2631–2637.

[63] Slattery M, et al. (2011) Cofactor binding evokes latent differences in DNA binding specificity between Hox proteins. Cell 147(6):1270–1282.

[64] (2015) UniProt: a hub for protein information. Nucleic Acids Research 43(D1):D204–D212.

[65] Maglott D, Ostell J, Pruitt KD, Tatusova T (2005) Entrez Gene: gene-centered information at NCBI. Nucleic Acids Research 33(suppl_1):D54–D58.

[66] Delaglio F, et al. (1995) NMRPipe: A Multidimensional Spectral Processing System Based on UNIX Pipes. Journal of Biomolecular NMR 6(3):277–293.

[67] Skinner SP, et al. (2015) Structure Calculation, Refinement and Validation Using CcpNmr Analysis. Acta Crystallographica Section D: Biological Crystallography 71(1):154–161.

[68] Liu Y, Schmidt B, Maskell DL (2010) MSAProbs: Multiple Sequence Alignment Based on Pair Hidden Markov Models and Partition Function Posterior Probabilities. Bioinformatics 26(16):1958–1964.

[69] Frishman D, Argos P (1995) Knowledge-Based Protein Secondary Structure Assignment. Proteins: Structure, Function, and Bioinformatics 23(4):566–579.

[70] Hubbard SJ, Thornton JM (1993) Naccess. Computer Program, Department of Biochemistry and Molecular Biology, University College London 2(1).

[71] Groussin M, et al. (2015) Toward More Accurate Ancestral Protein Genotype-Phenotype Reconstructions with the Use of Species Tree-Aware Gene Trees. Molecular Biology and Evolution 32(1):13–22.

[72] Jones DT, Taylor WR, Thornton JM (1992) The Rapid Generation of Mutation Data Matrices from Protein Sequences. Bioinformatics 8(3):275–282.

[73] Le SQ, Gascuel O (2008) An Improved General Amino Acid Replacement Matrix. Molecular Biology and Evolution 25(7):1307–1320.

[74] Yang Z (2007) PAML 4: Phylogenetic Analysis by Maximum Likelihood. Molecular Biology and Evolution 24(8):1586–1591.

[75] Edgar RC (2004) MUSCLE: multiple sequence alignment with high accuracy and high throughput. Nucleic Acids Research 32(5):1792–1797.

[76] Guindon S, et al. (2010) New algorithms and methods to estimate maximum-likelihood phylogenies: assessing the performance of PhyML 3.0. Systematic Biology 59(3):307–321.

[77] Otterbein LR, Kordowska J, Witte-Hoffmann C, Wang CLA, Dominguez R (2002) Crystal Structures of S100a6 in the Ca2+-Free and Ca2+-Bound States: The Calcium Sensor Mechanism of S100 Proteins Revealed at Atomic Resolution. Structure 10(4):557–567.

